# On the importance of time scales when studying adaptive evolution

**DOI:** 10.1101/269175

**Authors:** Charles Perrier, Anne Charmantier

## Abstract

Long-term field studies coupled with quantitative genomics offer a powerful means to understand the genetic bases underlying quantitative traits and their evolutionary changes. However, analyzing and interpreting the time scales at which adaptive evolution occurs is challenging. First, while evolution is predictable in the short term, with strikingly rapid phenotypic changes in data series, it remains unpredictable in the long term. Second, while the temporal dynamics of some loci with large effect on phenotypic variation and fitness have been characterized, this task can be complicated in cases of highly polygenic trait architecture implicating numerous small effect size loci, or when statistical tests are sensitive to the heterogeneity of some key characteristics of the genome, like recombination rate variations. After introducing these aforementioned challenges, we discuss a recent investigation of the genomic architecture and spatio-temporal variation in great tit bill length, which was related to the recent use of bird feeders. We discuss how this case study illustrates the importance of considering different temporal scales and evolutionary mechanisms both while analyzing trait temporal trends and when searching for and interpreting the signals of putative genomic footprints of selection. More generally this commentary discusses interesting challenges for unraveling the time scale at which adaptive traits evolve and their genomic bases.

**Impact summary:** An important goal in evolutionary biology is to understand how individual traits evolve, leading to fascinating variations in time and space. Long-term field studies have been crucial in trying to understand the timing, extent, and ecological determinants of such trait variation in wild populations. In this context, recent genomic tools can be used to look for the genetic bases underlying such trait variation and can provide clues on the nature and timing of their evolution. However, the analysis and the interpretation of the time scales at which evolution occurs remain challenging. First, analyzing long-term data series can be tricky; short-term changes are highly predictable whereas long-term evolution is much less predictable. A second difficult task is to study the architecture of complex quantitative traits and to decipher the timing and roles of the several genomic mechanisms involved in their evolution. This commentary introduces these challenges and discusses a recent investigation of the nature and timing of ecological and genomic factors responsible for variation in great tit bill length. Overall, we raise cautionary warnings regarding several conceptual and technical features and limitations when coupling analyses of long-term and genomic data to study trait evolution in wild populations.

## Main Text

Longitudinal field studies have brought invaluable insight for the understanding of evolutionary processes (Clutton-Brock & Sheldon 2010). Long-term studies notably allowed characterization of the temporality of trait variation and the strength and directionality of natural selection underlying such variation. Some of these long-term examinations of key phenotypic traits detected strikingly fast phenotypic change, driven by rapid ecological changes (Grant & Grant 2006). In contrast, many long-term studies failed to reveal micro-evolutionary change and response to selection (Merila *et al.* 2001; Pujol *et al.* 2018). While examining longitudinal data looking for both long-term trends as well as short-term fluctuations has the potential to shed light on evolutionary trajectories in natural populations, our ability to understand and more notably predict evolution remains limited outside the laboratory. Quantitative genetic models were initially developed with the aim of predicting evolutionary change, based on estimates of selection and additive genetic variation (Falconer 1960). Their predictive power worked efficiently for the genetic improvement of complex traits in many animal and plant breeding programs. Yet when spatio-temporal ecological heterogeneity is involved, evolution in the wild remains largely unpredictable (Pemberton 2010; Pujol et al. 2018).

Coupling such long-term studies with genomic tools is a powerful way to improve our understanding of the genetic bases underlying evolutionary changes in response to environmental variation. Rapid and recent monogenic adaptations based on de-novo mutations are often used as examples, for instance the rise of the melanic morph of the peppered moth *Biston betularia* following the industrial revolution (van’t Hof et al. 2016). Similarly, there are famous examples of rapid parallel monogenic or oligogenic adaptation based on long lasting standing genetic variation segregating in heterogeneous environments, for example coloration in deer mice *Peromyscus* and armor plates in stickleback *Gasterosteus aculeatus* (Barret & Hoekstra 2011; Nelson & Cresko 2018). However, these genomic analyses are facing several challenges when it comes to making inferences about polygenic adaptation and quantitative trait evolution. First, loci effect sizes are often small, requiring thousands, if not millions, of both SNPs and individuals for genome wide association studies (GWAS) to reveal significant effects. Such an investigation requires technical commitments (Wellenreuther & Hansson 2016; Gienapp et al. 2017a), but also a conceptual shift towards suppressing our desire to discover large effect alleles that are, in theory, rarely responsible for quantitative variation (Rockman 2011). Second, genome characteristics and especially variation in recombination rate along the genome can cause variation in the extent of background selection (defined as the loss of genetic diversity at a neutral locus due to negative selection against linked deleterious alleles) (Charlesworth et al. 1993; Charlesworth 2012; Nordborg et al. 2009) that can confound or bias detections of positive selection and hence our comprehension of the timing and nature of evolutionary trajectories based on genomic data (Roesti et al. 2012; Burri et al. 2015; Berner & Roesti 2017; Burri 2017; Comeron 2017; Delmore et al. 2018). Nevertheless, several studies have begun to decipher the polygenic mechanisms of rapid evolution of quantitative traits.

Avian bill morphology has played a prominent role in empirical studies of evolution and natural selection (Lack 1947; Grant 1999), perhaps because the size and shape of bills show large variations across and within bird species, and are shaped by strong selective forces since they directly determine foraging efficiency on various food sources. For instance, the emblematic study of Darwin’s finches on the Galapagos island of Daphne Mayor aimed at capturing evolutionary changes in bill size (Boag & Grant 1981; Grant & Grant 1993). From 1977 to 1978, bill size increased markedly after a severe drought in 1977. This analysis clearly demonstrated that extreme climatic events such as an *El niño* event are strong drivers of bill size evolution in this species. After 30 years of perspective, however, Grant & Grant (2002) concluded that while evolution of bill length was predictable as a rapid response to strong selection, it remained unpredictable on a slightly longer microevolutionary scale. Although the question of predictability of evolution across time scales remains challenging, even in the genomic era (Nosil et al 2018), genomic tools did provide insights on the evolution of bill size in the context of the rapid diversification of Darwin’s finches. A handful of genes were found to be significantly associated with bill size and shape in the medium ground finch *Geospiza fortis* (Lamichhaney et al. 2016), among which a major locus has been shown to influence bill dimensions in the Darwin finches’ entire radiation (Lamichhaney et al. 2015). These genomic analyses hence cracked the genomic architecture of this trait variation at both small and large microevolutionary scale, with the predominant control of a few large effect loci.

In a recent study, Bosse et al. (2017) investigated the genetic architecture of bill length in the Great tit *Parus major*, using a tremendous amount of data from long-term research programs in Wytham woods in the UK and in the Netherlands (NL). Applying modern analyses of both population and quantitative genomics using 500k single nucleotide polymorphisms, the authors provide insight into the signatures of divergent selection in the studied populations and the genomic architecture of variation in bill length. In line with the quantitative nature of variation in bill length and with quantitative genetics theory (Lynch & Walsh 1998), the authors showed that the genetic architecture of bill length was highly polygenic. Specifically, the authors showed, using a mixture analysis fitting all the SNPs simultaneously, that 3009 SNPs explained collectively 31% of bill length phenotypic variation. None of these SNPs reached genome wide significance in the GWAS with bill length, revealing small effect sizes of individual variants. In accordance with recent quantitative genomic findings notably for other traits in great tits (*eg* Robinson et al. 2013), the proportion of variance in bill length explained by each chromosome amazingly scaled with its size, demonstrating that the many SNPs additively explaining bill length are distributed throughout the genome. The polygenic analysis also predicted the difference in bill length observed between the UK and the NL, further illustrating the polygenic nature of this trait variation. This evidence for a polygenic control of bill length with no large effect SNPs is hence very different from the previous example on Darwin finches’ bill with large effect loci. Nevertheless, Bosse et al. (2017) also showed that variation at a single gene, col4A5, was associated with bill length. Bosse et al. (2017) then discussed whether an extended use of feeders, that are more abundant in the UK, might have driven the evolution of larger bills in the UK compared to the NL. Among the arguments pointing to feeders as drivers of longer bill lengths in UK great tits, Bosse et al. (2017) reported that col4A5 was associated with bill length in the UK but not in the NL, was highly differentiated between the UK and the NL, and was associated with greater reproductive success and higher activity at feeding sites in the UK. In addition, they reported that bills were longer in the UK compared to mainland Europe (Figure 4A in Bosse et al. 2017) and increased from 1982 to 2007 in the UK (Figure 4B in Bosse et al. 2017).

We argue in this comment that the speculation on the role of feeder on bill length evolution is not well supported by these arguments. Based on both phenotypic and genomic data, we propose instead that differences in bill length between the UK and the NL might have been evolving on a longer time scale than the contemporary use of feeders. First, an examination of bill length monitoring shows puzzling temporal and geographic patterns that may not incriminate the use of feeders. Second, the genomic patterns found at the region containing col4A5 might be compatible with the combined effect of variation in recombination rate and background selection, hence questioning the putative recent role of feeders as agents of positive selection.

The bill length trend inferred from the long-term monitoring is highly dependent on the time scale considered. Inspired by the readings of papers such as the Grant & Grant (2006) study relating the effects of rare and extreme events on the evolution of phenotypic traits, we carefully inspected the evolution of bill length during the studied period, looking for particularly rapid changes. We reanalyzed the data using a breakpoint computations method (coin R-package) aiming at localizing such striking change. The best cutpoint was found between 1986 and 1987 (maxT = 7.41, p-value < 2.2E-16), which corresponds to a conspicuous change in bill length. Measures from the five years preceding 1987 differed from subsequent years (t-test: t = −7.28, df = 674.53, p-value = 9.1E-13). Although a linear regression through the entire period, from 1982 to 2007, as implemented by Bosse et al., yielded a positive slope, removing the five measures prior to 1987, bill length significantly decreased from 1987 to 2007 (Figure 1A, Linear model from 1987 to 2007 either taking tarsus length into account, or not, F = 7.13, p-value = 8.2E-4; F = 13.2, p-value: 2.9E-4, respectively). A LOESS model confirms the decrease in bill length during the second part of the record. More generally, slopes of linear models linking bill length to time, using all of the possible combinations of 10 to 25 consecutive years (simply removing 1 to 16 years at the beginning, or the end, or both, of the data-series), were often positive when including one or more data points collected between 1982 and 1987, while negative slopes were often observed when excluding these years (Figure 2), further illustrating that both time periods yield opposite patterns. Furthermore, tarsus length also decreased significantly both over the 1987-2007 and the 1982-2007 periods (Figure 1B, F = 6.61, p-value = 0.01 & F = 24.07, p-value = 9.9E-07, respectively). Correspondingly, the bill length / tarsus length ratio did not change significantly from 1987 to 2007 (Figure 1C, F = 1.51, p-value = 0.28), indicating no change in bill length, when scaled to tarsus length, over this period. The origin of this cutpoint is important to investigate, since it could constitute an evidence for a response, either genetic or plastic, of both bill and tarsus lengths to a rapid abiotic or biotic change or to a methodological change. Yet, it seems at present difficult to decipher whether bill length or tarsus length or both, if any, were targeted by selection. Indeed, both traits are commonly phenotypically and genetically positively correlated in passerines (Teplitsky et al. 2014; Poissant et al. 2016), hence should often evolve together. Overall, these results very likely rule out the possibility of a contemporary (1982-2007) positive effect of feeders on bill length.

**Figure 1.**
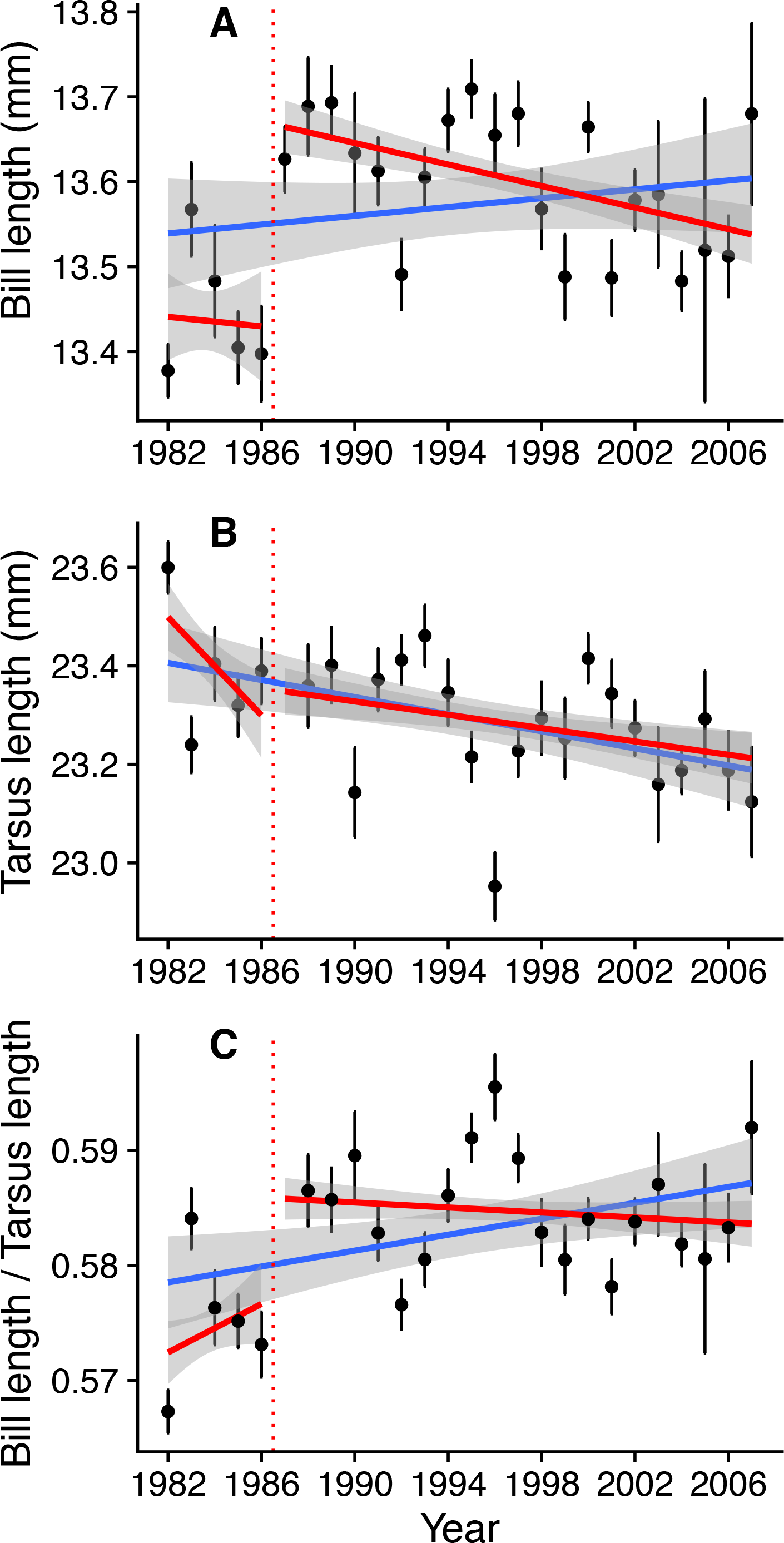
Decrease of mean bill length (A) and tarsus length (B) but conservation of the allometry between both traits (C), from 1987 to 2007 (red lines) in Wytham great tits. Regressions from 1982 to 2007 as in the original paper by Bosse et al. are shown by the blue lines. Dots and bars illustrate means and standard errors, respectively, for each year for each variable. Shaded areas represent 95% standard error around regressions lines.

**Figure 2.**
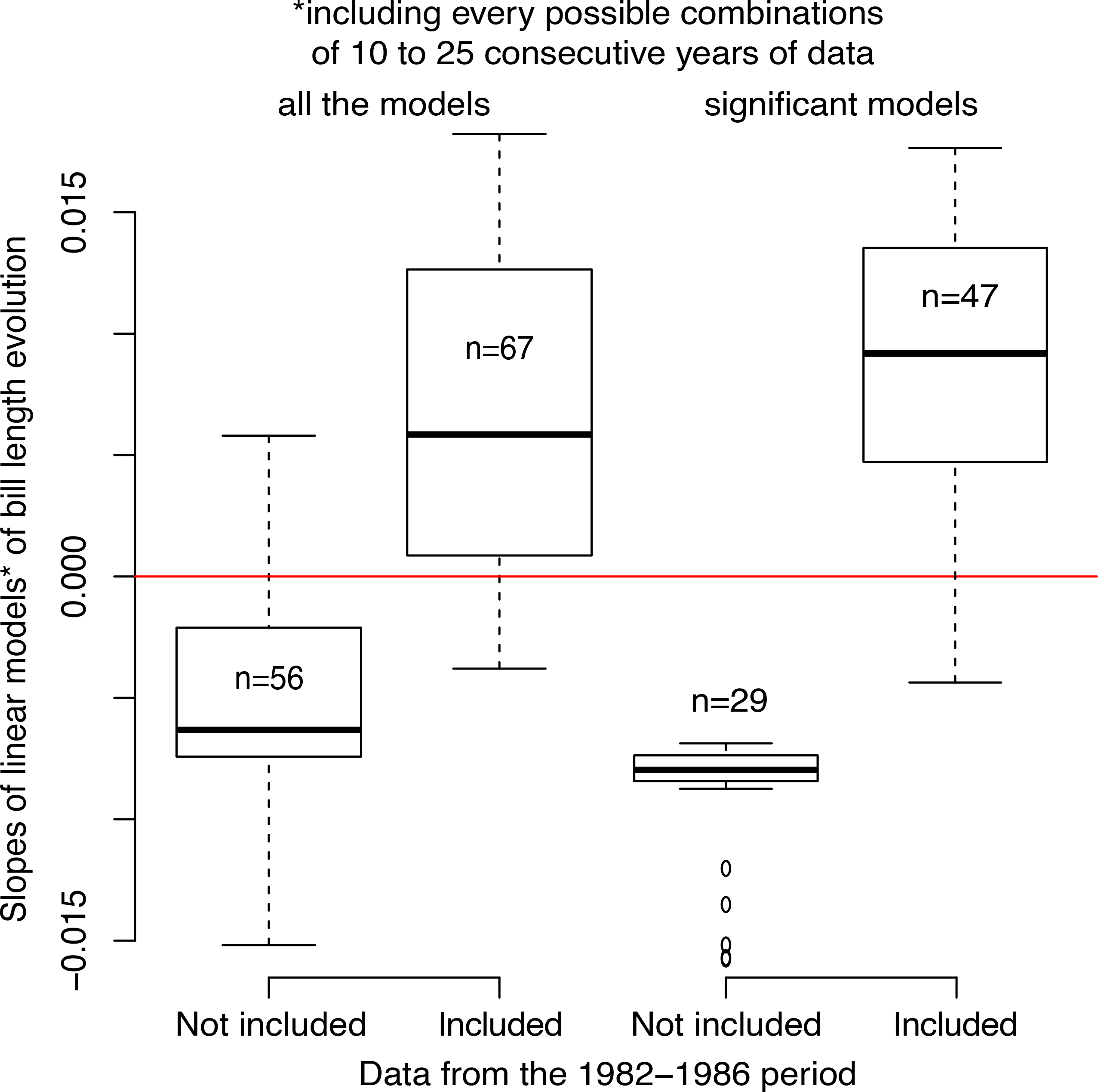
Slopes of linear models of bill length temporal evolution, including every possible combination of 10 to 25 consecutive years of data, with or without one to several years of data from 1982 to 1986.

One could nevertheless argue that bill length could have increased at a longer time scale than the 1982-2007 period, but still recent enough to incriminate the use of feeders. Figure 3 however shows the absence of a clear pattern of bill length increase in museum specimens collected in the UK. Although more data are needed to confirm the pattern, it suggests that larger bill length in the UK seems to have evolved over a longer time period than the one during which feeders have been used. Moreover, bill length across Europe does not display a clear dichotomy between the UK and mainland Europe but rather smooth spatial variations (Figure 4), with an ANOVA showing a significant effect of country (p = 3.75e-07) but not of sex (p = 0.345), with UK birds indeed having longer bills compared to three other countries (France, Italy, and the NL). This spatial variation also challenges the suggestion by Bosse et al. (2017) of a recent increase in bill length in British great tits caused by feeders.

**Figure 3.**
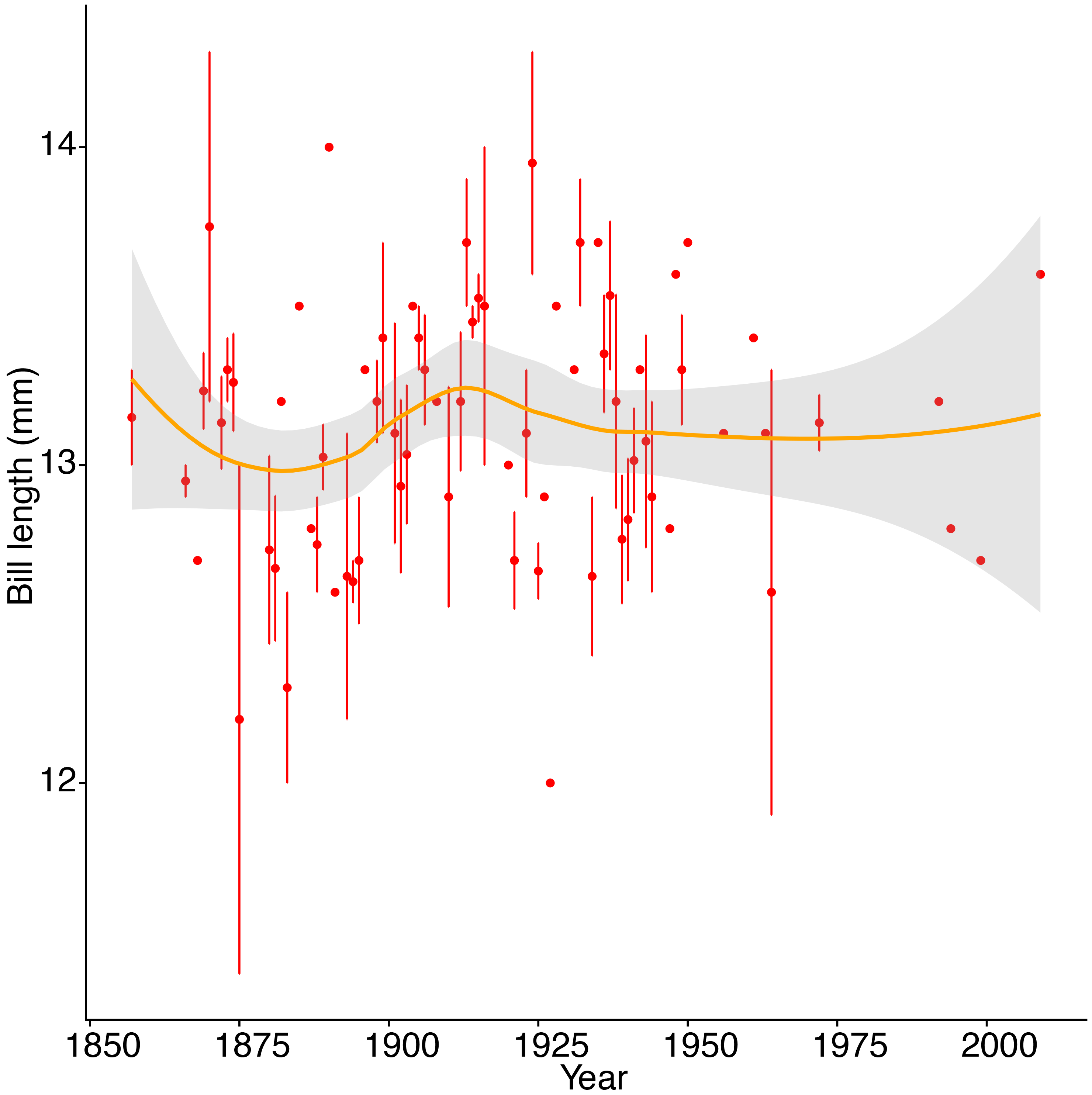
Variation in mean bill length in the UK from 1850 to 2007, based on museum records used in Bosse et al. 2017.

**Figure 4.**
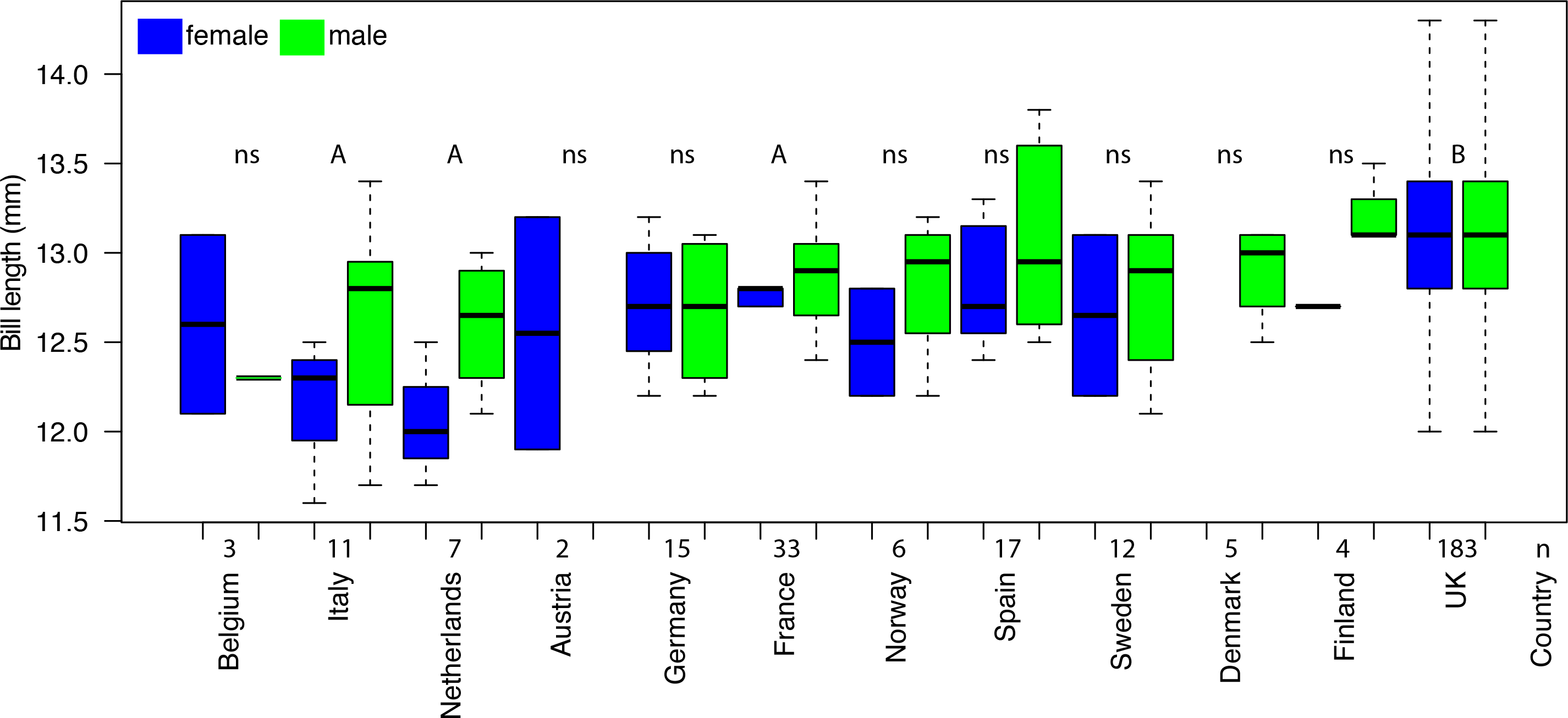
Variation in bill length across European countries for both female (in blue) and male (in green) great tit specimens from museums. Letters A and B illustrate significant differences between sites and ‘ns’ refers to non-significant differences (Tukey HSD).

We then question whether the evolution of the genomic region containing the gene col4A5, which was the corner stone linking bill length, activity at feeders, reproductive success and divergent selection, could have been influenced by recombination rate variation and background selection, rather than recent positive selection due to feeders. While the decay of LD is very fast in the great tit genome (marginal after 2kb, as shown in Figure S1 in Bosse et al. 2017, and in Figure 5A here (see supplementary material 1 for method details)), col4A5 was found in a large (>1Mb) genomic region with high long distance (20 to 200Kb) LD in the UK (Figure 5B). The F_ST_ between the UK and the NL was high along this region (Figure 5D here; figure S3A in Bosse et al. 2017). The eigenGWAS in this region was significant using both populations simultaneously for the entire set of SNPs (upper panel figure S2 in Bosse et al. 2017) and for chromosome 4A only (Figure 5E). As argued by Bosse et al. (2017), such results are compatible with recent strong positive selection in the UK over this large stretch of DNA on chromosome 4A. However, this large stretch of high LD around col4A5 was not only found in the UK but also in the NL (Figure 5C). The eigenGWAS in this region was also significant for both populations separately (Figure 5F & G). Furthermore, this large region was previously identified by Laine et al. (2016) as showing a signature of selective sweeps and reduced nucleotide diversity at the scale of the entire species distribution and not only in the UK (Sixth chromosome in Figure 2 in Laine et al. 2016). This same region was also identified as showing elevated differentiation in several lineages and populations of *Ficedula* flycatchers (Ellegren et al. 2012; Figure 1C in Burri et al. 2015). Therefore, given the existence of this large genomic region with reduced variation, increased LD, and increased divergence at several spatial scales in great tits, and in flycatchers, it seems unlikely that the mechanism shaping this region has been acting only in the UK, recently, and implying mainly positive section. Burri et al. (2015) determined that the high differentiation and high LD at this region, shared across flycatcher lineages, was due to the effect of linked selection combined with low recombination and issued a crucial warning: "scans are likely to identify recombination-mediated elevations of differentiation not necessarily attributable to selective sweeps". Accordingly, we propose that low recombination (potentially reflecting pericentromeric regions) and background selection in the region containing col4A5 in great tits could have resulted in locally reduced genetic diversity, increased differentiation and increased LD that could have altogether mimicked signatures of recent positive selection.

**Figure 5.**
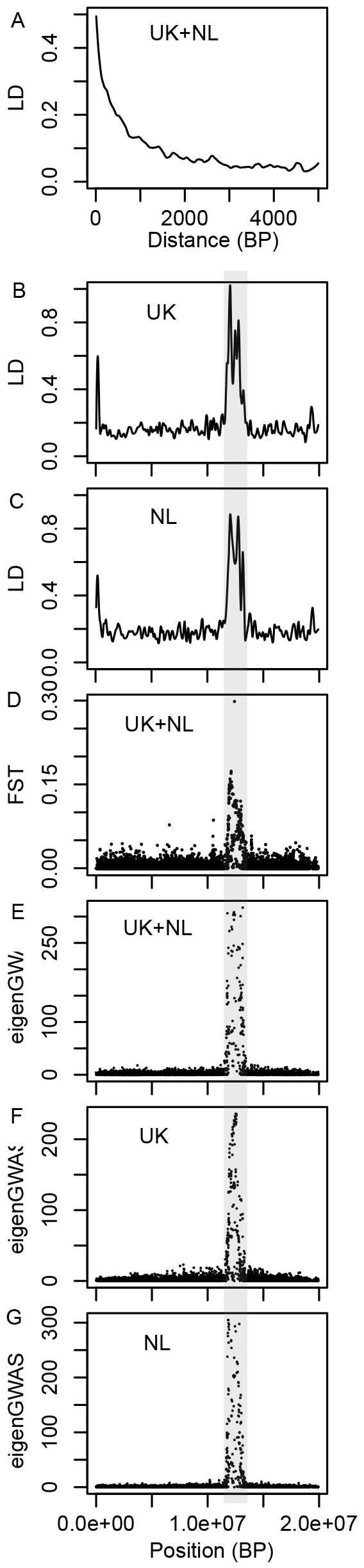
Genomic re-analysis of the chromosome 4A using the data from Bosse et al (2017). A) Smoothed linkage disequilibrium (LD) decay with genomic distance in UK and NL (pooled) great tits; Smoothed long distance (between SNPs distant from 20 to 200Kb) LD variation in UK (B) and NL (C); D) Single-marker F_ST_ between UK and NL; Single-marker significance of eigenGWAS for chromosome A4 in the entire sample (E) and only UK (F) and NL (G). The grey area indicates the region in which col4A5 is located.

To illustrate the general challenge raised here in differentiating recombination rate variation combined with background selection from positive selection, we present a very simple simulation of a 20Mb chromosome (hence comparable to the length of the chromosome 4A) containing a region of 1.6Mb with greatly reduced recombination (comparable to the length of the col4A5 haplotypes), with random occurrence of neutral and deleterious mutations but no beneficial ones (see supplementary material 2 for methods details). We show that for an average LD decay comparable to what was found in great tits (Figure 6A), long distance LD (Figure 6B), eigenGWAS (Figure 6C), integrated extended haplotype homozygosity (Figure 6D) and nucleotide diversity (Figure 6E) all show striking deviations in the recombination coldspot compared to the rest of the chromosome. Therefore, this simple simulation illustrates that such a pattern as that observed by Bosse et al. (2017) is compatible with the combined action of large-scale variation in recombination and background selection.

**Figure 6.**
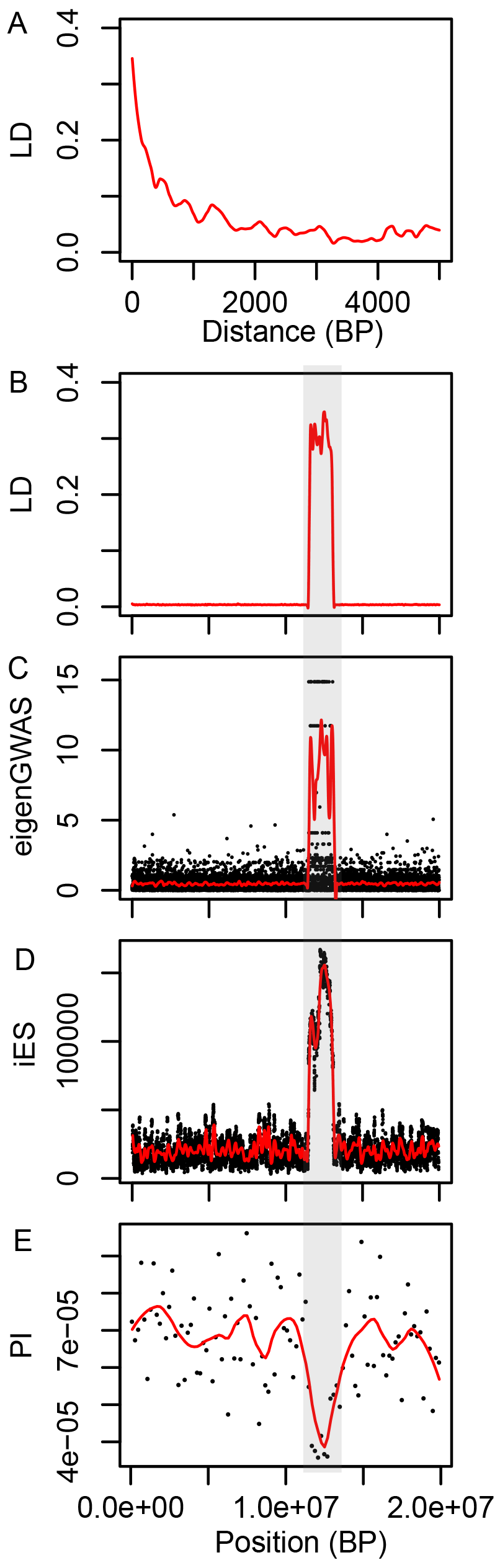
Genomic analysis of a 20Mb chromosome evolving in a single population, with a 1.6Mb region of reduced recombination (indicated by a grey area), and random occurrence of neutral and deleterious mutations and absence of beneficial mutations, simulated in SLiM. A) Smoothed LD decay with genomic distance, B) Smoothed long distance (between sites distant from 20 to 200Kb) LD variation, C) Single-marker and smoothed significativity of eigenGWAS, D) Single-marker and smoothed integrated Extended Haplotype Homozygosity (iES), E) 200kb and smoothed nucleotide diversity.

Neglecting the effects of variation in recombination and LD along the genome might have not only resulted in false positive footprints of selection in coldspots of recombination but also in false negatives in regions with a higher recombination rate (Berner & Roesti 2017). The large sliding window used in Bosse et al. (sliding over 200kb while the average LD is highly reduced after 2Kb) potentially worsened this problem by capturing principally elevated differentiation in large regions with increased LD possibly resulting from extended lower recombination due to structural variations. It may have also diluted narrow (*ie* narrower than the sliding window) peaks of elevated differentiation in more common regions with lower LD outside of recombination coldspots. Unfortunately, this probably relatively common caveat holds regardless of the nature of selection (*ie* positive or purifying) and is a purely mathematical problem of mismatch between the sliding window length and the extent of LD variation along the genome. It typically occurs when the unit of the sliding window is the base pair instead of the centimorgan. In fact, such large sliding windows relative to the LD decay should be used as neutral local baselines to ascertain the effects of variation in LD and recombination along the genome, with local outliers detected when comparing local residuals to such baselines (Roesti 2012; Burri 2017).

All these considerations suggest that, although the pattern of increased divergence at col4A5 is apparently compatible with strong positive selection, as suggested by Bosse et al (2017), the combined role of background selection, strong recombination rate variation and invariably large averaging window while long distance LD is variable, should be more comprehensively tested. Correspondingly, including these factors may also uncover more variants under stronger positive selection located outside of the few low recombination regions. In this context, the causality of the associations between col4A5 and bill length, activity at feeders and reproductive success in the UK, requires further clarifications.

This case study offers exciting avenues of research to unravel the determinants of both recent and long-term as well as spatial variations in quantitative traits in the Great tit but also in other emblematic species displaying quantitative trait variations. Spatial trait variation could be unraveled in multi-trait GWAS using polygenic frameworks, inspired by a pioneering study on great tit breeding phenology (Gienapp et al. 2017b). Additionally, a formal selection analysis relating bill length to over-winter survival when birds are most likely to benefit from food provided by people, is required to elucidate the nature of the evolutionary forces behind bill length variation. Finally, including the effect of variation in recombination rate, background selection and LD along the genome to draw local neutral envelopes of genomic differentiation and modulate the local width of sliding windows, will likely help identify further candidate loci (Roesti et al. 2012; 2013; Burri 2017; Comeron 2017; Delmore et al. 2018). Genomic analyses performed in other populations across Europe could also help determine the timing and the nature of selection, by taking into account the temporal dynamics of the differentiation landscape (Burri 2017). We wish to conclude by emphasizing the importance of integrating, or at least recognizing, the widespread and sometimes strong variation in recombination rate along genomes, which can in some circumstances distort our understanding of evolutionary processes based on genomic investigations (Roesti et al. 2012; Burri et al. 2015; Berner & Roesti 2017; Burri 2017; Comeron 2017; Delmore et al. 2018).

## Acknowledgements

We thank several colleagues from the Institute of Evolutionary Sciences of Montpellier (ISEM) and from the Center for Functional and Evolutionary Ecology (CEFE), especially Paul Jay, Tim Janicke, Nicolas Bierne, Pierre-Alexandre Gagnaire, François Bonhomme, and Patrice David, for discussions regarding the impact of recombination rate variation on the landscape of diversity and differentiation. We thank Caroline Rose for comments on the English syntax. We thank the associate editor, three anonymous reviewers and JC Senar for constructive comments on previous versions of the manuscript. Both authors are presently funded by a European Research Council Starting grant - *Selection in Heterogeneous Environment* (ERC-2013-StG-337365-SHE). The authors declare no conflict of interest.

Supplementary material 1. Reanalysis of the COL4A5 region. We downloaded the SNP dataset from Bosse et al. 2017 and used vcftools (Danecek et al. 2011) to estimate LD across the genome for the entire dataset. We then estimated LD for both populations separately along the chromosome 4A. To represent long distance LD, we kept LD values between each pairs of markers distant from 20kb to 200kb. We smoothed these statistics using the loess function native to R. We estimated F_ST_ using Plink (Purcell et al. 2007). We performed eigenGWAS tests (Chen et al. 2017) in the UK, the NL and both. EigenGWAS were performed in chromosome 4A only and not on the entire genome, in order to be more comparable with the simulation outputs on one chromosome. Results from whole genome eigenGWAS can be found in figure S2 in Bosse et al. (2017).

Supplementary material 2. Simulation of a chromosome similar in size to the chromosome 4A in great tit, and containing a region of reduced recombination of approximately the same size as the COL4A5 region. We used SLIM (Haller et al. 2017) to simulate a very simple scenario of reduced recombination in a 1.6Mb region within a 20Mb long chromosome. Recombination rate was equal to 1e-5 from 0 to 11.5Kb and from 13.1Kb to 20Mb. Recombination rate was equal to 1e-100 11.5Kb to 13.1Kb. Population size was equal to 1000. Simulation lasted 2000 generations. Mutation rate was le-7. We implemented 3 types of mutations: i) neutral, 50% of the mutations, ii) slightly deleterious (fitness effect of −0.1), 25% of the mutations, iii) severely deleterious (fitness effect of −0.9), 25% of the mutations. We exported a VCF dataset containing about 8500 SNPs from a randomly chosen simulation. We used vcftools to estimate LD across the chromosomes and nucleotide diversity (PI) over 200 kb windows. To represent long distance LD, we kept LD values between each pairs of markers distant from 20kb to 200kb. We performed eigenGWAS tests (Chen et al. 2017). We estimated iES (Sabesti et al. 2007) using the R package rehh (Gautier et al. 2016). We smoothed statistics using the loess function native to R.

## References and Notes

Barrett RD & Hoekstra HE (2011). Molecular spandrels: tests of adaptation at the genetic level. Nature Reviews Genetics, 12, 767.

Berner D, Roesti M (2017) Genomics of adaptive divergence with chromosome-scale heterogeneity in crossover rate. Molecular Ecology, 26, 6351–6369.

Boag PT, Grant PR (1981) Intense natural selection in a population of Darwin’s finches (Geospizinae) in the Galapagos. Science, 214, 82–85.

Bosse M, Spurgin LG, Laine VN et al. (2017) Recent natural selection causes adaptive evolution of an avian polygenic trait. Science (New York, N.Y.), 358, 365–368.

Burri R (2017) Interpreting differentiation landscapes in the light of long-term linked selection. Evolution Letters, 1, 118–131.

Burri R, Nater A, Kawakami T et al. (2015) Linked selection and recombination rate variation drive the evolution of the genomic landscape of differentiation across the speciation continuum of Ficedulaflycatchers. Genome Research, 25, 1656–1665.

Charlesworth B. (2013). Background selection 20 years on: the Wilhelmine E. Key 2012 invitational lecture. Journal of Heredity, 104(2), 161–171.

Charlesworth B., Morgan, M. T., & Charlesworth, D. (1993). The effect of deleterious mutations on neutral molecular variation. Genetics, 134(4), 1289–1303.

Clutton-Brock T, Sheldon BC (2010) Individuals and populations: the role of long-term, individual-based studies of animals in ecology and evolutionary biology. Trends in Ecology & Evolution, 25, 562–573.

Comeron JM (2017) Background selection as null hypothesis in population genomics: insights and challenges from Drosophila studies. Philosophical Transactions of the Royal Society B: Biological Sciences, 372, 20160471–13.

Delmore KE, Lugo Ramos JS, Van Doren BM et al. (2018) Comparative analysis examining patterns of genomic differentiation across multiple episodes of population divergence in birds. Evolution Letters, 2, 76–87.

Ellegren H, Smeds L, Burri R et al. (2012) The genomic landscape of species divergence in Ficedula flycatchers. Nature, 491, 756–760.

Falconer, D. S. (1960). Introduction to quantitative genetics. Oliver And Boyd; Edinburgh; London.

Gienapp P, Fior S, Guillaume F et al. (2017a) Genomic Quantitative Genetics to Study Evolution in the Wild. Trends in Ecology & Evolution, 1–12.

Gienapp P, Laine VN, Mateman AC, Van Oers K, Visser ME (2017b) Environment-Dependent Genotype-Phenotype Associations in Avian Breeding Time. Frontiers in Genetics, 8, 1392–9.

Grant BR, Grant PR (1993) Evolution of Darwin’s Finches Caused by a Rare Climatic Event. Proceedings of the Royal Society B-Biological Sciences, 251, 111–117

Grant PR (1999) Ecology and evolution of Darwin’s finches. Princeton University Press.

Grant PR, Grant BR (2002) Unpredictable Evolution in a 30-Year Study of Darwin’s Finches. Science, 296, 707–711.

Grant PR, Grant BR (2006) Evolution of character displacement in Darwin’s finches. Science (New York, N.Y.), 313, 224–226.

Haller BC & Messer PW (2017) SLiM 2: Flexible, interactive forward genetic simulations. Molecular Biology and Evolution 34, 230–240.

Lack D (1947) Darwin’s finches. Cambridge University Press, London.

Laine VN, Gossmann TI, Schachtschneider KM et al. (2016) Evolutionary signals of selection on cognition from the great tit genome and methylome. Nature Communications, 7, 1–9.

Lamichhaney S, Berglund J, Almén MS et al. (2015) Evolution of Darwin’s finches and their beaks revealed by genome sequencing. 1–16.

Lamichhaney S, Han F, Berglund J, Wang C, 2016 A beak size locus in Darwin’s finches facilitated character displacement during a drought, science.sciencemag.org

Lynch M, Walsh B (1998) Genetics and Analysis of Quantitative Traits. Sinauer Associates Incorporated.

Merila J, Sheldon BC, Kruuk LE (2001) Explaining stasis: microevolutionary studies in natural populations. Genetica, 112–113, 199–222.

Nelson TC & Cresko WA (2018). Ancient genomic variation underlies repeated ecological adaptation in young stickleback populations. Evolution Letters, 2, 9–21.

Nordborg M, Charlesworth B, & Charlesworth D (1996). The effect of recombination on background selection. Genetics Research, 67, 159–174.

Poissant J, Morrissey MB, Gosler AG, Slate J, Sheldon BC (2016) Multivariate selection and intersexual genetic constraints in a wild bird population. Journal of Evolutionary Biology, 29, 2022–2035.

Pemberton JM (2010) Evolution of quantitative traits in the wild: mind the ecology. Philosophical Transactions of the Royal Society of London B: Biological Sciences, 365, 2431–2438.

Pujol B, Blanchet S, Charmantier A et al. (2018) The Missing Response to Selection in the Wild. Trends in Ecology & Evolution, 1–10.

Rockman MV (2011) The QTN program and the alleles that matter for evolution: all that’s gold does not glitter. Evolution, 66, 1–17.

Roesti M, Hendry AP, Salzburger W, Berner D (2012) Genome divergence during evolutionary diversification as revealed in replicate lake-stream stickleback population pairs. Molecular Ecology, 21, 2852–2862.

Roesti M, Moser D, Berner D (2013) Recombination in the threespine stickleback genome-patterns and consequences. Molecular Ecology, 22, 3014–3027.

Teplitsky C, Tarka M, Møller AP et al. (2014) Assessing Multivariate Constraints to Evolution across Ten Long-Term Avian Studies. 9, e90444–15.

van)t Hof AE, Campagne P, Rigden DJ, Yung CJ, Lingley J, Quail MA […] & Saccheri IJ (2016). The industrial melanism mutation in British peppered moths is a transposable element. Nature, 534, 102.

Wellenreuther M, Hansson B (2016) Detecting Polygenic Evolution: Problems, Pitfalls, and Promises. Trends in Genetics, 32, 155–164.

